# Altered co-expression patterns of synovial fluid proteins related to the immune system and extracellular matrix organization in late stage OA, compared to non-OA controls

**DOI:** 10.1101/2023.03.20.533133

**Authors:** Jenny Lönsjö, Martin Rydén, Aleksandra Turkiewicz, Velocity Hughes, Jon Tjörnstand, Patrik Önnerfjord, Martin Englund, Neserin Ali

## Abstract

**Objective:** Synovial fluid contains proteins that may have been released from surrounding tissues, our aim was to gain new insights into the proteomic profiles of human synovial fluid in knees with and without osteoarthritis (OA).

**Methods:** We used synovial fluid from 11 patients with end-stage medial compartment knee OA, aspirated during total knee replacement, and from 13 deceased donors who had no prior history of knee OA (healthy controls). These samples were analyzed using high-multiplex immunoassays Olink®. The differential expression of proteins between the groups was analyzed using a linear mixed effects model. The linear associations between pairs of protein expressions were estimated with a linear regression model.

**Results:** We found that almost half of the detected proteins were differentially expressed between the OA and non-OA controls. The proteins that were most elevated in the OA group compared to controls were tartrate-resistant acid phosphatase type 5 (fold change 10.6, 95% CI [6.6-17.0]), coagulation factor XI (4.3 [2.6-6.8]) and urokinase-type plasminogen activator (4.3 [2.3-6.8]). The proteins with lower levels in OA compared to controls were fatty acid-binding protein, adipocyte (0.03 [0.02-0.05]), myocilin (0.05 [0.03-0.08]) and carbonic anhydrase 3 (0.14 [0.09-0.23]). The protein-protein co-expression analysis suggests an overall lower number of protein pairs that show co-expression in OA.

**Conclusion:** There is a substantial change in protein abundance in synovial fluid in end-stage knee OA, suggesting that global joint homeostasis is severely deranged. Our findings suggest altered co-expression between the immune response and extracellular matrix organization in end-stage knee OA, in comparison to non-OA controls.

## Introduction

Despite ongoing research, knowledge of the pathogenesis of osteoarthritis (OA) at the proteomic level is still sparse and inconclusive (1). This lack of knowledge hampers the development of methods for reliable diagnosis, as well as further advancements regarding treatments (1). OA is a multifactorial disease that affects the whole joint, through biomechanical, metabolic, and immune response pathways (2). The disease is primarily characterized by bone remodeling, degradation of cartilage, and synovial inflammation. Drugs targeting these pathogenic processes are currently being evaluated(3). However, they have not been proven to be effective thus far. This strongly indicates that further efforts are needed to understand the complex molecular pathways.

Synovial fluid (SF) is in direct contact with most knee tissues involved in the OA process, such as the hyaline cartilage, menisci, and synovium. Thus, SF reflects the molecular environment of the joint, and therefore represents an ideal fluid to examine patterns of molecular change during OA. Proteins serve essential functions in this environment, including enzymatic reactions to activate or deactivate specific metabolic pathways, immunological responses or cellular signals. The dynamic status of the proteome makes comprehensive analysis of synovial fluid more challenging. However, in addition to identification and quantification of individual proteins, understanding and characterizing the dynamic crosstalk between proteins present in a sample is essential for comprehensive understanding of the underlying biology. New insights into potential differences in the proteomic characteristics of SF in healthy knees and knees with OA could, for instance, pave the way for better understanding of OA pathogenesis, detection of new biomarkers, and for the identification of new treatment strategies in OA.

Explorative proteomics is commonly performed by liquid-chromatography mass-spectrometry (LC-MS), due to its ability to identify over 1500 sample specific proteins (in synovial fluid) in an un-targeted manner. However, targeted antibody-based assays such as from Olink Proteomics® (Uppsala, Sweden) can contribute to new insights, providing complementary knowledge regarding the proteomic composition of SF in health and disease. The main advantage of the Olink assay is its ability to detect low-abundance proteins, which are often masked by high abundance proteins when analyzed with MS. Another advantage is the ability to analyze crude samples, avoiding the sample preparation steps needed prior to MS-analysis (4, 5).

The aims of our study are: 1) to identify differentially expressed proteins in synovial fluid from knees with or without OA and 2) to investigate if the patterns of protein-protein co-expression are different between the two groups. As there is lack of coverage on proteomic profiles of SF from human knees analyzed by multiplex assay panels, we do not specify an *a priori* hypothesis about differences in expression. We consider this to be an exploratory study.

## Materials and methods

### Sample selection

The synovial fluid samples used in this study came from patients who had total knee replacement (TKR) due to end-stage medial compartment knee OA (Outerbridge grade 4) and from deceased donors without known knee OA. The OA-group was made up of 11 individuals, 8 women and 3 men between 55-80 years of age, and the healthy group was made up of 13 individuals, 8 women and 5 men between 19-79 years of age. Further information regarding the patients and donors can be found in Supplementary Table 1. All tissue samples collected from deceased donors were required to be retrieved within 48 hours post-mortem and stored in −80°C within 2 hours of retrieval. A prerequisite for a knee to be considered healthy was that the menisci were macroscopically intact and that there was no visible damage on the articular cartilage surface from the medial and lateral compartment. In addition, subjects with visible blood contamination in the SF were not included. In the OA group, SF was collected by transcutaneous aspiration prior to the TKR procedure. SF-samples from both groups were centrifuged (1800 rpm for 10 min), and the supernatant was separated from the pellet prior to being frozen. The supernatant was used for this study. The study was approved by the ethical review committee of Lund University (Dnrs: 2015/39;2016/865; 2019/3239).

### Sample preparation and Olink analysis

The SF samples were thawed and diluted 1:2 with MilliQ water. 10 µl of each diluted sample were pipetted into a 96 well plate and sealed, prior to being sent for analysis at Olink. The samples were analyzed by Olink Proteomics, using proximity extension assays (PEA) (6, 7). For each specific protein assay, two antibodies targeting different epitopes of the same protein are coupled to complementary DNA strands. The physical proximity of the antibodies when bound to their respective epitopes enables their complementary DNA strands to adhere to each other and get replicated. The specificity of the antibodies, as well as the need for proximity for the matching DNA strands to anneal, provides a protein-specific detection method. Each DNA strand is uniquely barcoded for its corresponding protein and sample, enabling a multiplex assay. The double stranded DNA segments are then amplified through real-time PCR, quantifying the amounts of different proteins in the sample. Results from the OLINK assay are obtained as Normalized Protein eXpression (NPX (log_2_ scale)) values for each protein, which can be related to protein concentration (pg/ml). We employed three panels: the Cardiometabolic (CAM), Cardiovascular III (CVD III) and Development (DEV) panels, each testing for 92 proteins. The samples were run at the Olink facility. Due to possible saturation problems, additional runs were performed with additional dilutions of the samples: 1:200 and 1:1000 for CAM and 1:20 and 1:200 for CVD III. The DEV panel was not run at any additional dilutions.

### Statistical analysis

The NPX values were provided by Olink. To estimate the difference in protein expression between the groups, we used a hierarchical linear mixed effects model. The person was included as a random effect, while the protein, group (healthy vs OA) and their interactions were included as fixed effects. The model was further adjusted for age, sex and BMI. Hierarchical models allow for shrinkage of the variance, by borrowing information between proteins. All proteins were included in one model. All estimates of the between-group differences in expression of each protein were provided with nominal 95% confidence intervals (CI). Due to the use of one hierarchical model and the exploratory nature of the study, no multiplicity corrections were performed.

In total, three proteins were excluded from the analysis. Cartilage oligomeric matrix protein (COMP) and chitinase-3-like protein 1 (CHI3L1) were excluded based on the appearance of their dilution plots (see **Error! Reference source not found**. and **Error! Reference source not found**.). The relationship between the different dilutions was far from the predicted. In addition, it was decided to exclude proteins if more than two samples in the same group had values below the Limit of Detection (LOD). This was the case for von Willebrand factor (vWF), which was therefore also excluded.

To be able to compare the NPX values for a certain protein between samples, the same dilution needs to be used for all samples. Furthermore, for the measured NPX value to be reliable, it needs to be higher than the Limit of Detection (LOD) and lower than the Upper Limit of Quantification (ULOQ), making up the dynamic range of the assay. When more than one dilution was feasible to use, we used the dilution that gave an NPX value that was closest to the middle of the dynamic range, based on the appearance of the dilution plot (**Error! Reference source not found**. and **Error! Reference source not found**.). How this was performed is further explained in the supplementary data. The chosen dilutions for all included proteins are presented in **Error! Reference source not found**..

### Protein-protein co-expression

Linear associations between the expression levels of pairs of proteins were estimating by fitting a linear regression model with the expression of one of the proteins as outcome, and the expression of the other protein, the group, and their interaction as independent variables. The NPX values were standardized by subtracting the mean protein expression and dividing by the standard deviation, to make the slopes comparable between models.

### Pathway enrichment analysis

Pathway enrichment analysis was conducted using the Reactome database. Three different enrichment analyses were conducted: one for the proteins that were differentially expressed between the OA and non-OA group, and two separate ones for the proteins included in the protein-protein co-expression analysis: one for the healthy group and one for the OA group. To be included in the enrichment analyses for proteins from the protein-protein co-expression analyses, proteins were required to differ in co-expression slopes between OA and non-OA by ± 1 or more, and the base 2 log 95% CIs of the slopes were required not to cross over 0. The background proteome included all Olink proteins detected in this analysis.

### Differential network analysis

As an exploratory approach to more advanced co-expression analysis, we used undirected gaussian graphical models using joint graphical lasso approach (8). The obtained network shows conditionally dependent pairs of proteins, where the density and similarity of the networks are guided by selection of two tuning parameters. We have selected the tuning parameters attempting to obtains models that are easy enough to interpret, but complicated enough to be interesting, judging the model fit using AIC and BIC. We selected 3 models of increasing complexity based on these metrics. The results should be treated as hypothesis generating due to very exploratory nature of the method.

### Data visualization and presentation

The slope estimates that differed between non-OA and end-stage knee OA were further plotted in a protein-protein network, with the proteins as nodes and the estimated slope values as the edges between the nodes, using the igraph R package. The selected proteins co-expression data is presented in a network when the following criteria below were fulfilled a) The CI for the differences in slopes between OA and non-OA did not include 0 and a) the slope in non-OA was at least 1 (absolute value) and CI not spanning 0 or b) the slope in OA was at least 1 (absolute value) and CI not spanning 0.

### Reproducibility and quality control

No duplicates were run. However, more than one dilution was run for the CAM and CVD III panels. Dilution plots were created to assess if the dilutions were within the assay’s dynamic range, and if a greater or lower dilution was more appropriate. These plots can be found in the supplementary data. In addition, samples below the LOD were considered to be missing and were excluded. The assay manufacturer Olink Proteomics also performed a quality control (QC) for each sample to ensure that the analysis was performed in a correct manner considering the technical aspects.

## Results

### Differentially expressed proteins

The differential expressions of 274 proteins were compared between the end-stage OA and non-OA group and plotted (Supplementary Table 3). Out of these, 35 proteins are down-regulated, and 105 proteins are up-regulated in end-stage OA compared to non-OA controls, with 95% confidence intervals of the base 2 log fold-change not spanning zero. These proteins are displayed in Figure 1. Thirty-three of these proteins have a fold change of magnitude 1.5 or more, whereof 12 were downregulated**Error! Reference source not found**.. The most pronounced differences between end-stage OA and the controls were found for tartrate-resistant acid phosphatase type 5 (PPA5), which showed 10.6 fold increase with a 95% CI [6.6, 17.0], plasminogen activator inhibitor 1 (PA1) (5.0 [3.1, 8.0]), coagulation factor XI (F11) (4.3 [2.7, 6.9]), urokinase-type plasminogen activator (uPA) (4.3 [2.6, 6.8]), and C-type lectin domain family 11 member A (CLEC11A) (4.2 [2.6, 6.8]). The proteins with reduced levels in end-stage OA in comparison to controls were fatty acid-binding protein, adipocyte (FABP4) (0.03 [0.02, 0.05]), myocilin (MYOC) (0.05 [0.03, 0.08]), carbonic anhydrase 3 (CA3) (0.14 [0.09, 0.23]), and myoglobin (MB) (0.18 [0.11, 0.28]).

**Figure1.**
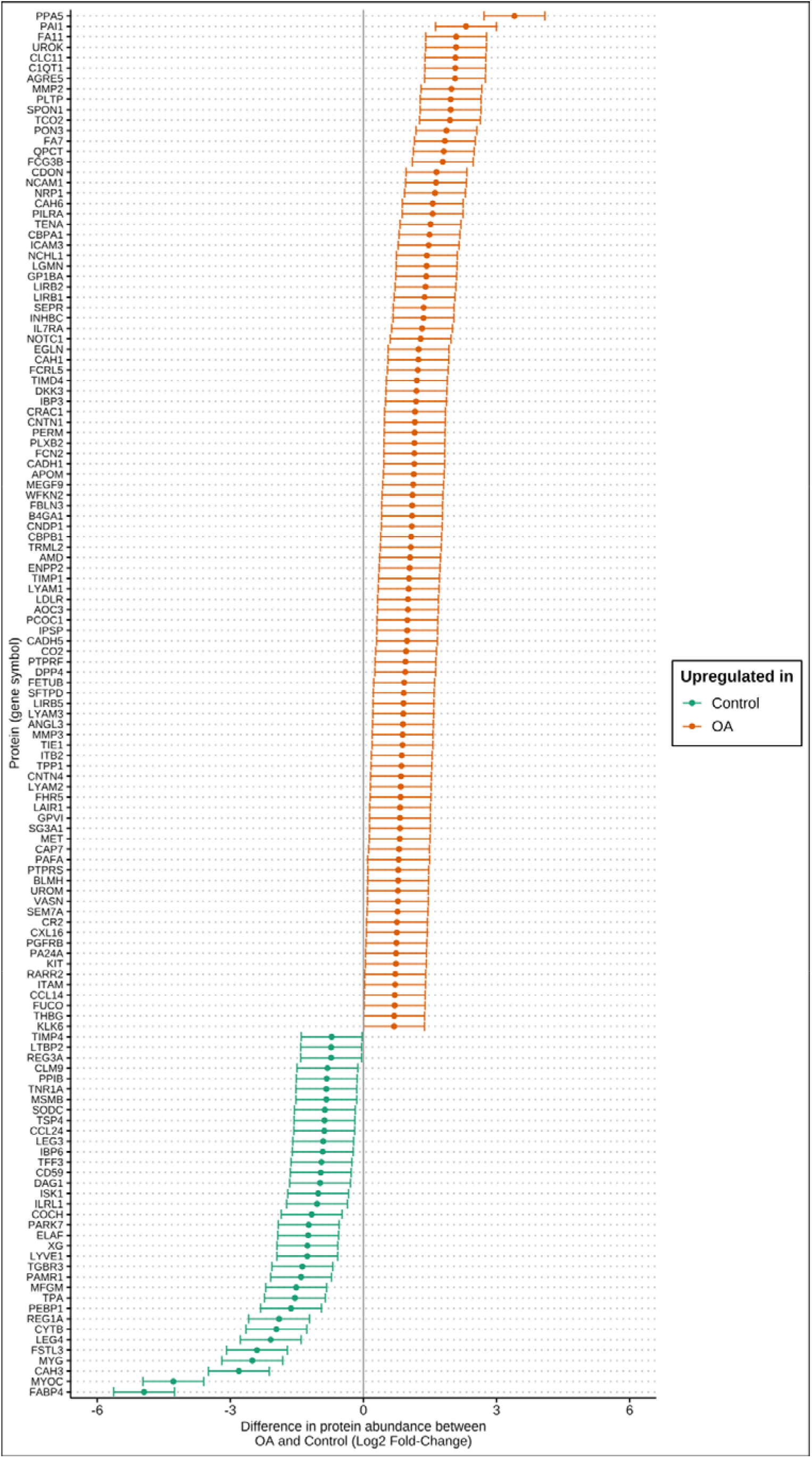
Differential expression for proteins confidence intervals (CI) of 95%.

### Protein-protein interaction networks

Overall, there appeared to be more protein pairs that showed positive co-expression rather than negative co-expression in both groups (Figure 2). The co-expression networks were highly dense and interconnected. Several proteins showed a difference in co-expression patterns between the OA and non-OA group (Figure 3, Supplementary Table 4). In non-OA SF, ACAN showed negative co-expression with IL18BP (slope: −0.86), ICAM1 (−0.83), SELP (−1.02), GUSB (−0.86), KIT (−0.81), and MMP9 (−1.0), but lost its co-expression in end-stage OA, while NOTCH3 was negatively co-expressed with ACAN in non-OA SF (−0.50) and positively co-expressed with ACAN in end-stage OA (0.90). In non-OA SF, TMSB10 was negatively co-expressed with SIRPB1 (−1.61), IL6RA (−1.68), IGFBP6 (−1.6), SERPINA7 (−1.86), and PROC (−1.76), but lost this negative co-expression in end-stage OA, while CR2 (−1.55) went from a negative co-expression in the non-OA donors to a positive co-expression in end-stage OA (0.54). In non-OA SF, VISG4 had many negative co-expressions with other proteins, with most of these co-expressions lost in end-stage OA. However, its negative co-expressions with CDH5 (−0.44), NOTCH1 (−0.43), CNTN1 (−0.42), CCL24 (−0.74), ICAM3 (−0.40), MEGF9 (−0.44), CR2 (−0.52), NCAM1(−0.50) and CHLI (−0.44) in non-OA SF became positively co-expressed in end-stage OA. CCL5 had a positive co-expression with CRIM1, KLK6, DLK-1. GP6, LTBR, CD300LG, CR2, NOTCH3, CD93, COL18A1, CD99L2, ANGPTL3, PEAR1, NOV, ESAM, CNTN4 and CD58, but showed negative co-expression with these in the end-stage OA group.

**Figure 2.**
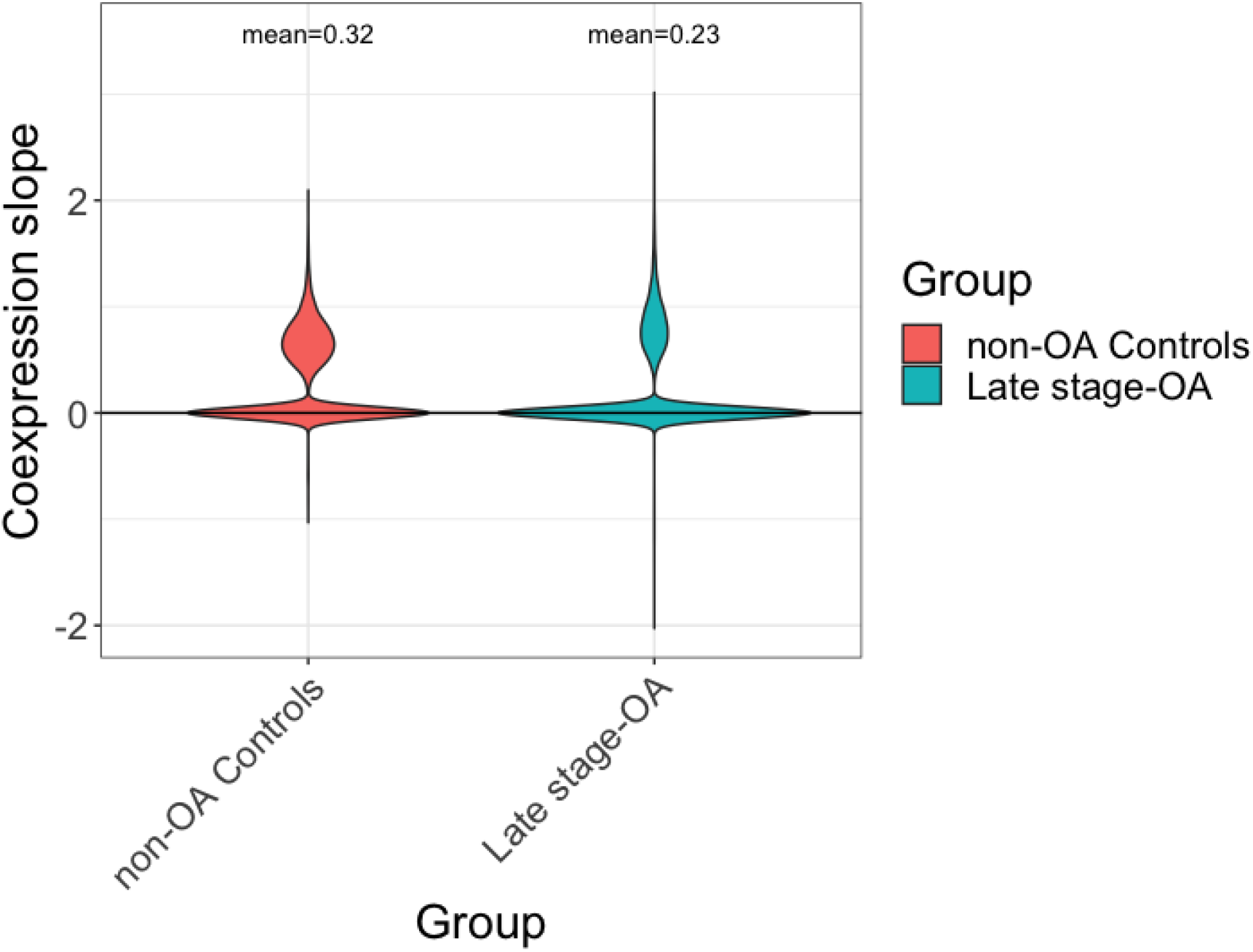
Estimated co-expression between proteins, “slope” represents the association between expression levels of protein pairs.

**Figure 3.**
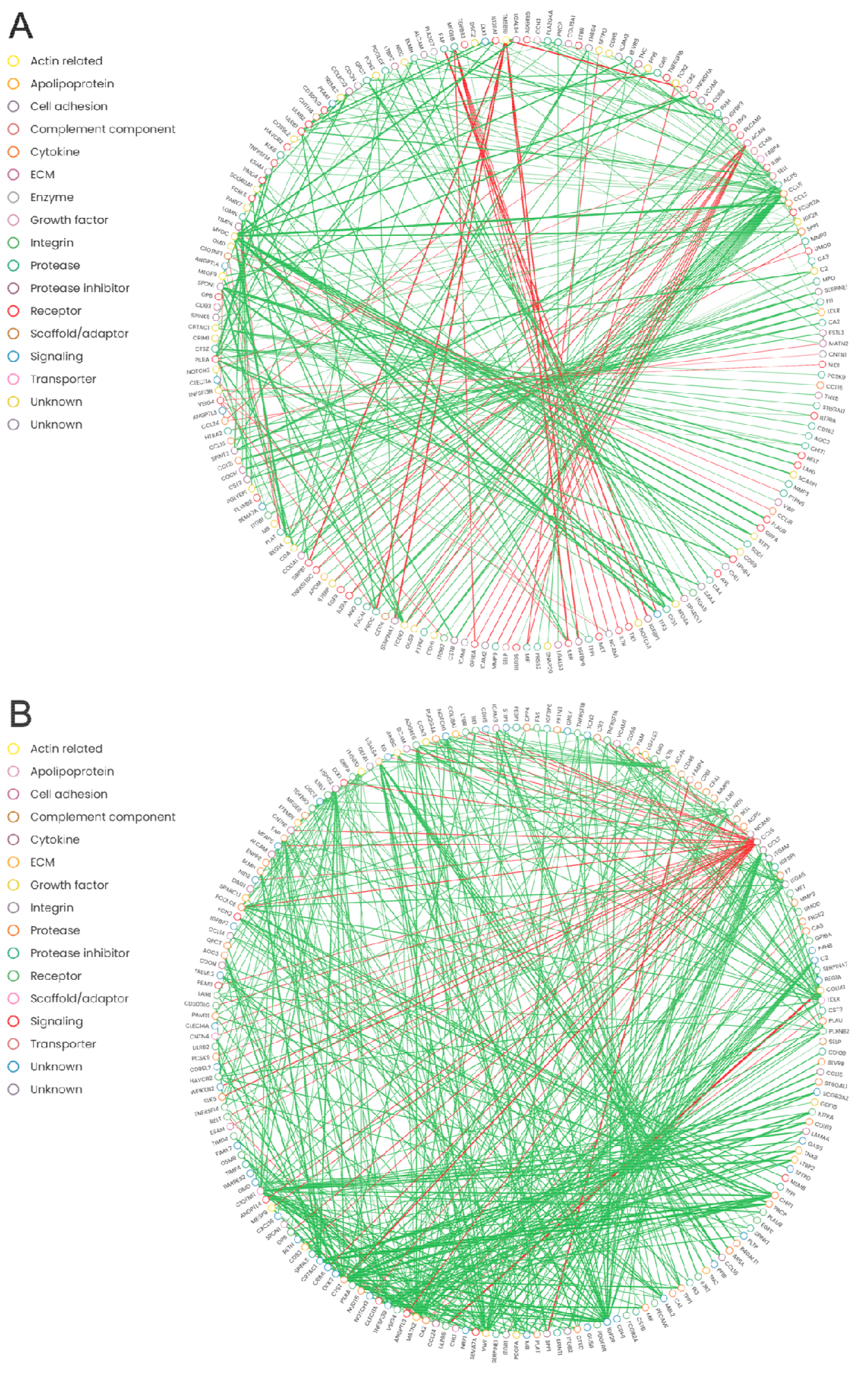
Protein-protein interaction networks based on the co-expression slopes. Only the co-expression slopes that differed with an absolute value of 1 or more between non-OA control and late stage OA are included with an addition of having a 95% confidence interval of the base 2 log fold-change not spanning over zero were used in the interaction network. These networks represent the co-expression slopes that were different between the groups. The nodes (proteins) are color coded based on their known molecular function. The red and green lines represent if it’s a negative=red or positive=green co-expression. A) non -OA controls co-expression B) OA co-expression

### Differential network analysis

The differential network analysis showed different clusters of proteins in the non-OA vs OA group. Clusters like NOTCH1-ICAM-CHLI-MEGF9, LILRB1-CNTN1 and IL18BP-CD46-IGLC2-LYVE1 were more prominent in non-OA SF and clusters like MCP1-ALCAM-SEMA7A-IGF2R, ENG-PLXNB2-ITGA5 and HAVCR2-TNFR1-SPRINT1 were more prominent in end-stage OA (Supplementary document).

### Reactome pathway enrichment analysis

Pathways that were enriched in the differentially expressed proteins were associated with signal transduction, the immune system, hemostasis and extracellular matrix organization (Table 1, Supplementary Table 5, 6 and 7). The same types of pathway associations were found for the protein lists from the protein-protein co-expression data. However, there was a clearer enrichment of extracellular matrix organization-associated proteins in the co-expression dataset, in comparison to the differentially expressed protein data set. Proteins that showed greater co-expression in non OA SF in comparison to end-stage OA showed an enriched immune system association. Conversely, extracellular matrix organization proteins and signal transduction pathways were more enriched in proteins that showed more co-expression in OA in comparison to the healthy.

## Discussion

Almost 50% of the detected proteins differed in abundance between the non-OA and end-stage medial compartment knee OA samples. If this result is representative of the majority of the proteins in synovial fluid (i.e. including the ones not tested in the Olink panels), this could suggest that much of the proteome is widely different in end-stage OA in comparison to a healthy state. Also, the protein-protein co-expression analysis showed a distinct difference in co-expression patterns in the end-stage OA vs the non-OA group, with fewer proteins overall showing co-expression in end-stage OA.

Twelve out of the 33 proteins that have an absolute log2 fold change (FC) of 1.5 or greater are down-regulated in the end-stage OA samples. Some of them have previously been mentioned to be associated with the progression of OA. Fatty acid binding protein 4 (FABP4) helps solubilize fatty acids to facilitate their transport intracellularly, and it has been proposed as a possible biomarker for OA. Previous studies have reported that downregulation of FABP4 promotes osteoclast formation (9). However, contrary to our findings, it has been reported that FABP4 is up-regulated in SF and plasma in individuals with OA (10, 11). Follistatin-like protein 3 (FSTL3) is a glycoprotein that is involved in regulating inflammation and suppresses osteoblast differentiation of bone marrow mesenchymal cells during that process(12). FSTL3 has been reported to be both down- and up-regulated in SF and serum, respectively, in samples from patients diagnosed with OA (13, 14). Previous findings regarding FABP4 and FSTL3 are somewhat contrasting to our results, and they need to be investigated further to be able to conclude if the development of OA always triggers a decrease in these proteins. Carbonic anhydrase 3 (CA3) helps maintain an appropriate pH level in cells by converting carbon dioxide to bicarbonate, they have been found to have a protecting activity for the osteocytes from oxidative stress. Osteocytes control osteoblast and osteoclast activities both directly via cell-to-cell communication and indirectly via secreted factors(15). CA3 has not previously been reported to affect the development of OA. However, auto-antibodies targeting CA3 have been reported to be involved in rheumatoid arthritis, contributing to a decrease in CA3 in serum samples of patients with this disease (16). Myocilin (MYOC) has previously been mentioned to be expressed at a lower level in healthy medial meniscus compared to OA medial meniscus (17). MYOC is a secreted glycoprotein that has been reported to be expressed in bone marrow-derived mesenchymal stem cells (MSCs) and plays a role in their differentiation into osteoblasts *in vitro* and in osteogenesis *in vivo*. The cortical bone thickness and trabecular volume a known factor of bone remodeling and osteoblast differentiation, were reduced dramatically in the femurs of Myoc-null mice compared with wild-type mice. It has been suggested that MYOC should be considered as a target for improving the bone regenerative potential of MSCs. The decreased differential expression promotes an increased osteoclast formation and decreased osteoblast formation.

18 proteins are up-regulated in the end-stage OA group compared to the non-OA group, with a fold change of 1.5 or more. Previously studies have mentioned some of them as possibly playing a role in the development of knee OA. The presence of neural cell adhesion molecule 1 (NCAM1), also referred to as CD56, has not been examined in SF before. However, it has been reported to be up-regulated in peripheral blood, and down-regulated in an experimental OA model based on chondrocytes (18, 19). Literature suggests that NCAM silencing inhibited osteoblastic differentiation (20), and that its presence can regulate chondrocyte hypertrophy in chromogenic differentiation (18). The protein Fc fragment of IgG receptor IIIb (FCGR3B) is involved in the immune response and are known to bind to IgG, and has been detected to be more abundant in menisci with OA than in healthy menisci (21). Spondin 1 (SPON1), also known as F-spondin, has a neuroregulatory function, and has been reported to be up-regulated in cartilage from individuals with OA compared to control specimens(22). In vitro, SPON1 has been shown to directly stimulate the differentiation and proliferation of osteoblasts and decrease osteoclast differentiation in periodontal tissue (23). In our study, the protein with the second highest fold-change in end-stage OA vs non-OA controls was plasminogen activator inhibitor 1 (SERAPINE1), which acts as an inhibitor of tissue plasminogen activator (tPA) and urokinase (uPA), which block the intrinsic pathway and coagulation(24). uPA is an enzyme responsible for the cleavage of plasminogen to form plasmin. Plasmin mediates the degradation of the extracellular matrix either by itself or in conjunction with matrix metalloproteinases. Matrix metallopeptidase 2 (MMP2) is an enzyme with proteolytic activity, and it has been stated that the level of MMP2 is elevated in SF from individuals with OA compared with healthy SF(25). We also observed an increase of both coagulation factor VII (F7) and XI (F11) in end-stage OA. Both of these proteins are involved in the intrinsic pathway of blood coagulation. Their role in the development of OA is still not clear, but recent studies have reported that there is constant crosstalk between the coagulation and inflammation pathway (26). The protein with the highest fold change were PPA5, it is a known osteoclasts-produced enzyme during bone resorption. In cell culture, its activity is used to detect and enumerate osteoclasts. CLC11, another protein with a very 4.2 log2fold change, is known to promote osteogenesis by stimulating the differentiation of mesenchymal progenitors into mature osteoblasts. The mechanisms that are presented by these proteins is a combination of enzymes that are responsible for the degradation or to set the right environment for the enzymatic activity to take place and proteins involved in both osteoclast and osteoblast activity. These are signs that both a degradation and degeneration is taking place in these joints reflected in the outcome of these activities in SF from knee OA patients.

Our exploratory co-expression and network analysis highlight clusters of proteins that are not necessarily differentially abundant between the groups, but could be interesting. The protein co-expression analysis suggests a negative association between ACAN and proteins like IL18BP in a healthy synovial fluid, while this protein is linked to, CD46, IGlC2 and LYVE1 in our exploratory network analysis. Previous studies have reported that ACAN, which is in high abundancy in normal cartilage, has a negative association with an increased severity of OA. Even though our confidence interval for difference in ACAN levels between OA and healthy was inconclusive, our previous mass spectrometry results from the same samples suggested a decrease of ACAN abundancy in late stage OA in comparison to healthy (27). These discrepancies could due to methodological differences, mass spectrometry searches for trypsin digested peptides while Olink targets specific epitopes. Additionally, ACAN is a 250kDa protein and different domains are known to be processed differently by various proteases during pathology. The biological association between ACAN and IL18BP is not clear, but IL-18BP have shown to suppress IL-17-induced osteoclastogenesis and rectifies T cell imbalance in rheumatoid arthritis(28). The cluster found in the differential network analysis is connecting IL-18BP to CD46 which is involved in the protective aspect of the immune system (29). None of these proteins showed a clear increase or decrease when comparing their abundancy between healthy and OA, on the other hand their protein-protein co-expression as well as the differential expression network was clearly different between the healthy condition vs OA, indicating an altered immune response communication with ECM proteins like ACAN.

VSIG4 is a strong negative regulator of T-cell proliferation, some of the proteins that showed a negative co-expression with VSIG4 in non-OA but a positive co-expression in end-stage OA showed to form a cluster in the differential network analysis, MEGF9-ICAM3-NOTCH1-CHLI. MEGF9 are thought to be involved in the development, maintenance and injury response of the nervous system. Upregulation of MEGF9 can increase the expression of EGFR, matrix metalloproteinase-13 and a disintegrin like and metallopeptidase with thrombospondin type 1 motif 5, so as to aggravate cartilage degradation. ICAM3 bind to the leukocyte adhesion LFA-1 protein. ICAM3 mediates adhesion between cells by binding to specific integrin receptors. It plays an important role in the immune cell response through its facilitation of interactions between T cells and dendritic cells, which allows for T cell activation. Comparative evaluation of leukocyte- and platelet-rich plasma and pure platelet-rich plasma aimed at cartilage regeneration suggested that the leucocyte rich platelet-rich plasma may be associated with less regeneration of cartilage due to a higher activation of NF-kB pathway (30). NOTCH1 is of the type 1 transmembrane protein family. Recently, the NOTCH pathway was identified as a potential regulator of both catabolic and anabolic molecules in the cartilage ECM during development(31-33). Interestingly, recent studies have also suggested that the NOTCH pathway is highly activated in mouse and human joint tissues during post-traumatic OA (25–27), and that temporary suppression of NOTCH signaling in murine joints leads to delayed OA progression (25). These data collectively suggest that physiological NOTCH signaling within joint tissues is essential for joint maintenance, however, when NOTCH signaling is abnormally activated such as during post-traumatic OA, temporary inhibition of the NOTCH pathway or its downstream effectors may provide a means for altering the progression of post-traumatic OA. Both MEGF9, NOTCH1 and CHLI was increased in OA. This network of proteins associated with degradation of ECM and infiltration of immune cells were found to be in the differential network of the healthy controls. This suggest that the increase of some of these proteins in OA might have affected their co-expression.

Immune response is increasingly considered the key factor affecting cartilage repair (34). It has both negative and positive regulatory effects on the process of regeneration and repair. Proinflammatory factors are secreted in large numbers, and necrotic cartilage is removed (34). Immune cells can secrete anti-inflammatory factors and chondrogenic cytokines, which can inhibit inflammation and promote cartilage repair (34). Our enrichment analysis suggests a higher ratio of proteins co-expressing, associated with both the immune system and the extracellular organization in healthy in comparison to OA. This could indicate an altered co-expression between the immune response and the extracellular degeneration.

Finally, we would like to discuss some limitations of our study. Because no previous study has evaluated the matrix effect of synovial fluid on the these specific Olink panels, there were challenges in determining the best dilution of the samples for optimal assay performance. For future Olink studies with SF, it might be suitable to optimize the appropriate dilution ratios with a few test samples prior to running all the study samples. Furthermore, in our study, the detected SF proteins are limited to the three chosen Olink panels. Thus, it would also be of interest to run SF samples using other panels in future experiments. Each panel targets a subset of proteins deemed to be of interest within a specific disease area, and despite some of these areas appearing to be quite distant from OA, only one protein out of 276 were excluded due to multiple values being below LOD. The sample size is relatively small, but the groups are well balanced with respect to age, sex and BMI, and we were also adjusted the analyses for those potential confounders. The co-expression and network analyses should be treated as exploratory, but we believe they could potentially raise interesting aspects of the data.

## Conclusions

We have identified a profound difference in the proteomic profiles of the synovial fluid from human subjects’ knees with end-stage knee OA vs no OA. It is evident that the progression of OA is reflected in the changes in levels of a broad subset of proteins. Our findings indicate that there still are several proteins that previously have not been reported to be involved in the pathogenesis of OA, and that there is a need to continue to investigate proteomic changes during the development of the disease to fully understand its pathogenesis and different endotypes.

