## Supplementary material for "Altered co-expression patterns of synovial fluid proteins related to the immune system and extracellular matrix organization in late stage OA, compared to non-OA controls": Supplemetary document

Neserin Ali

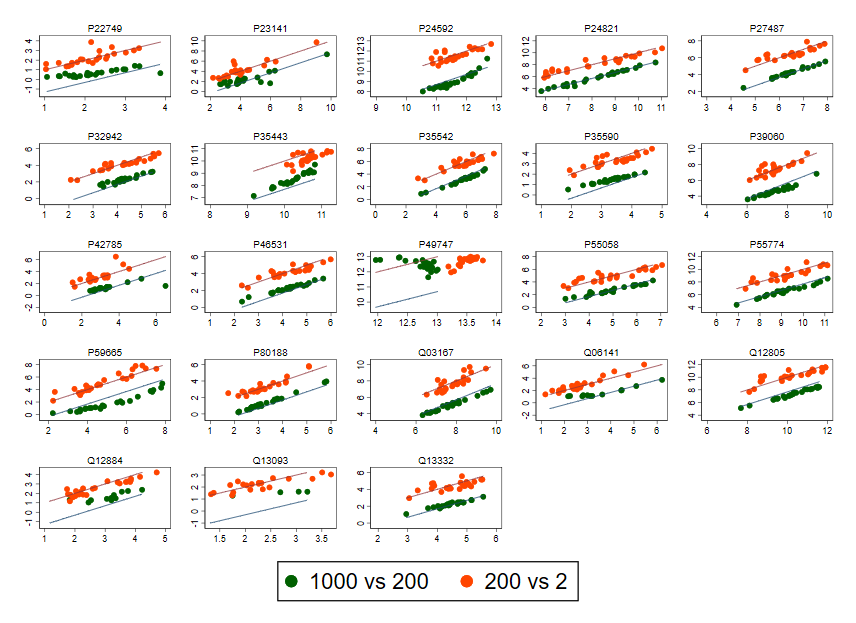

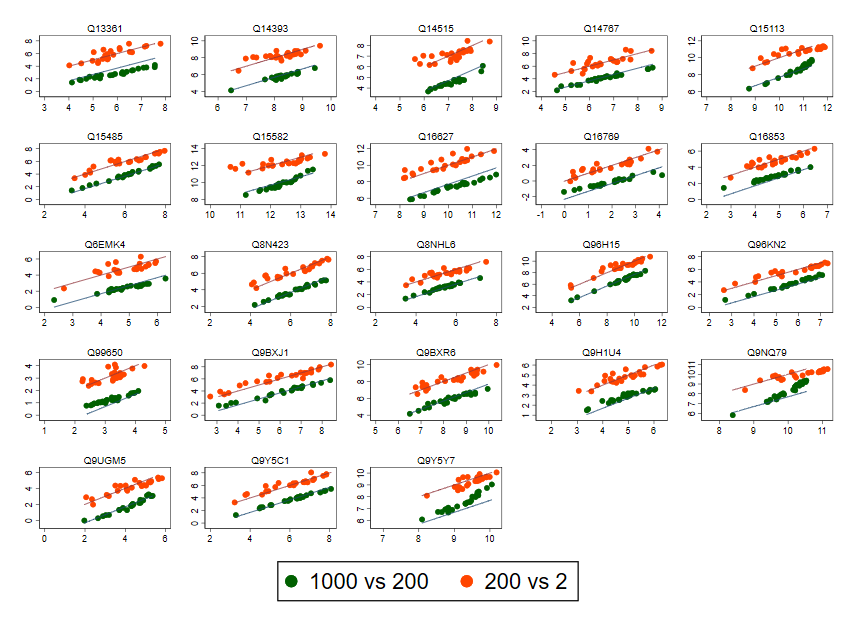

Continued on the next page.

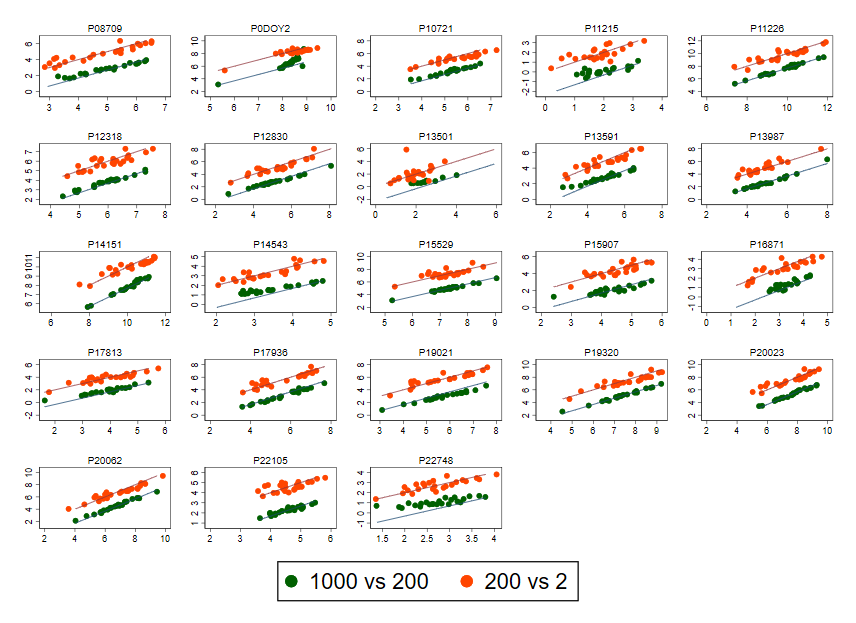

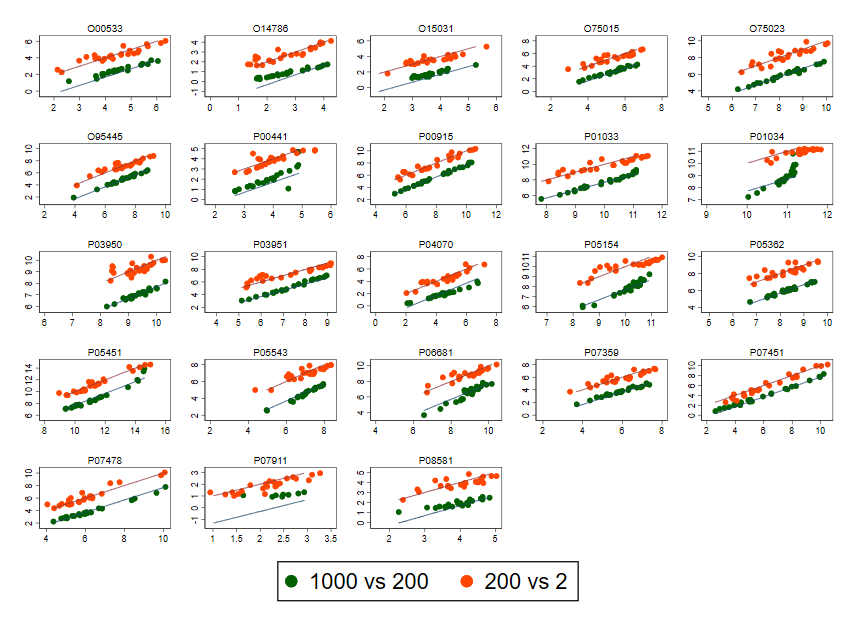

Supplementary figure 1.Dilution plots for all proteins in the CAM panel. The plots display the correlation between the different dilutions.

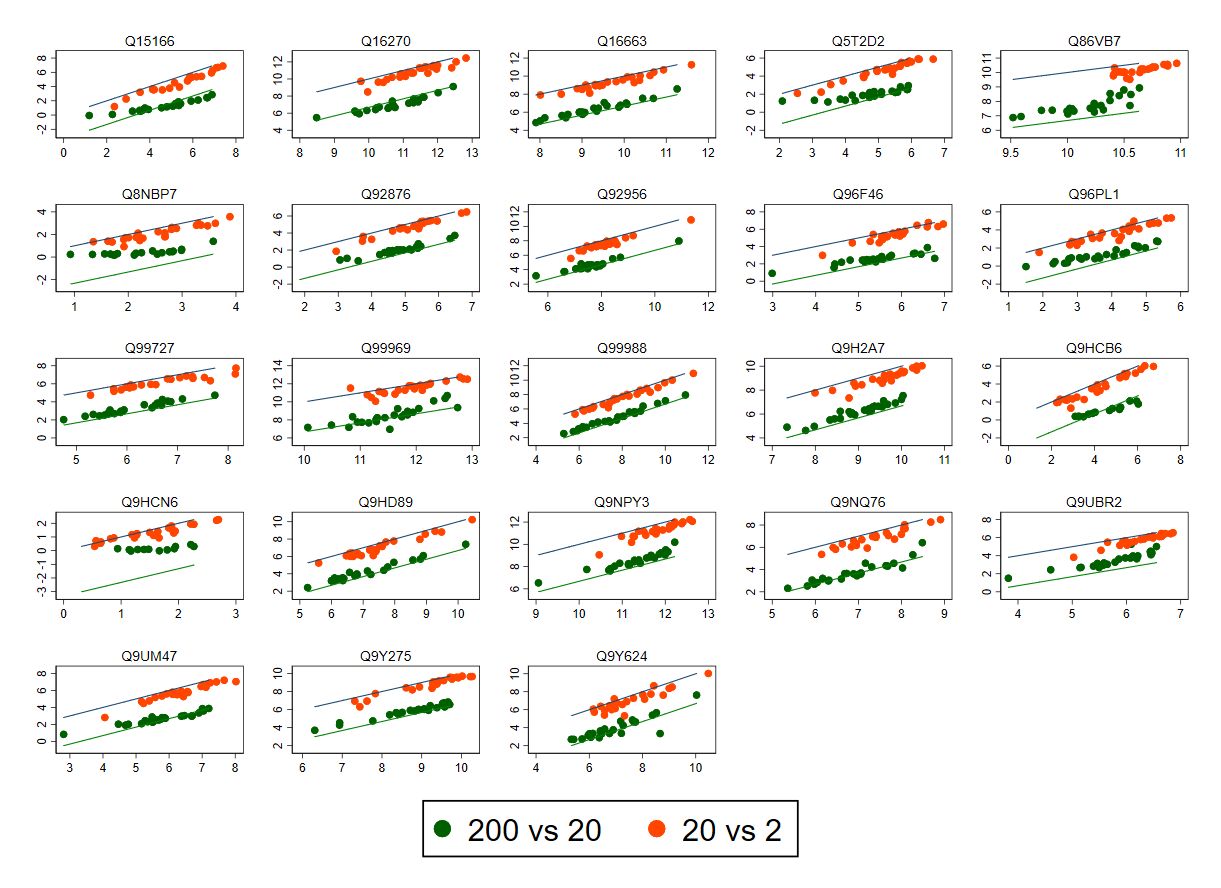

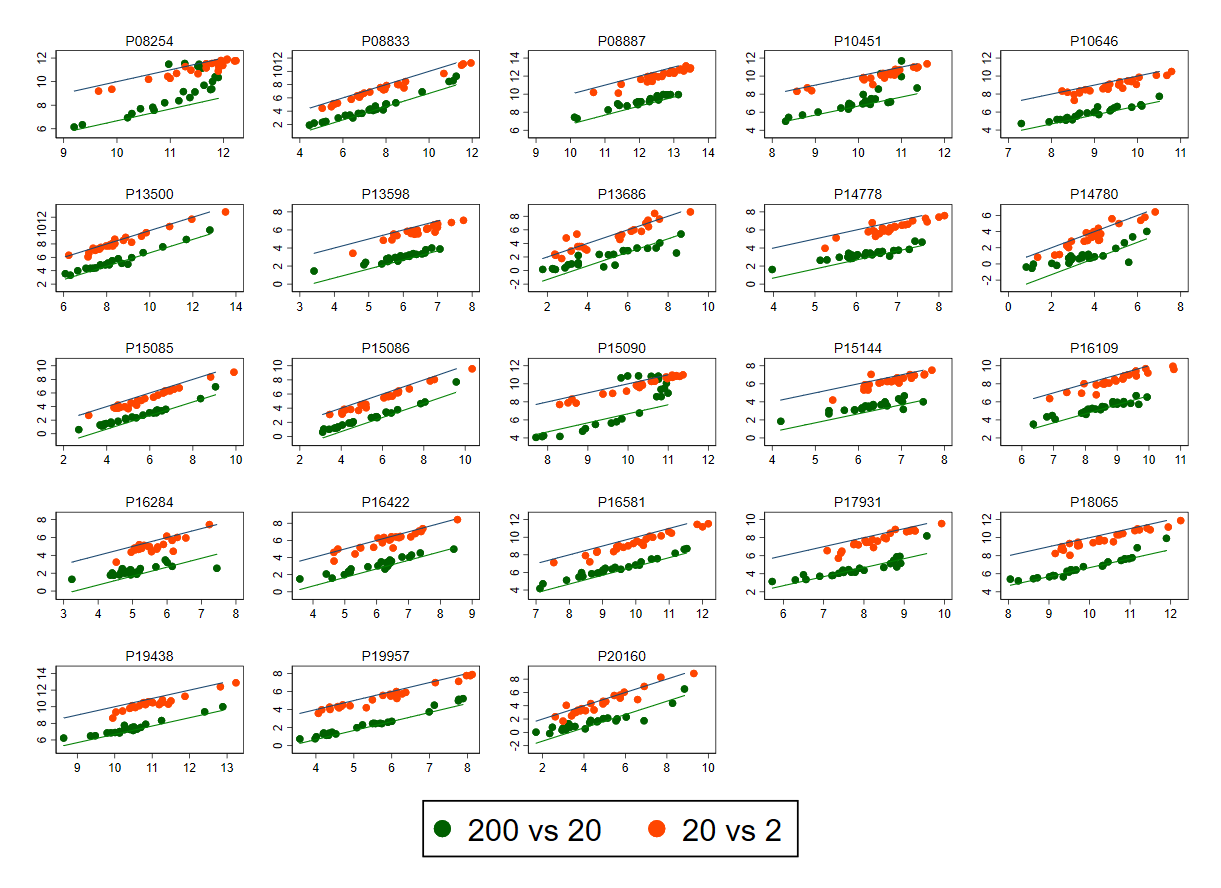

Continued on the next page.

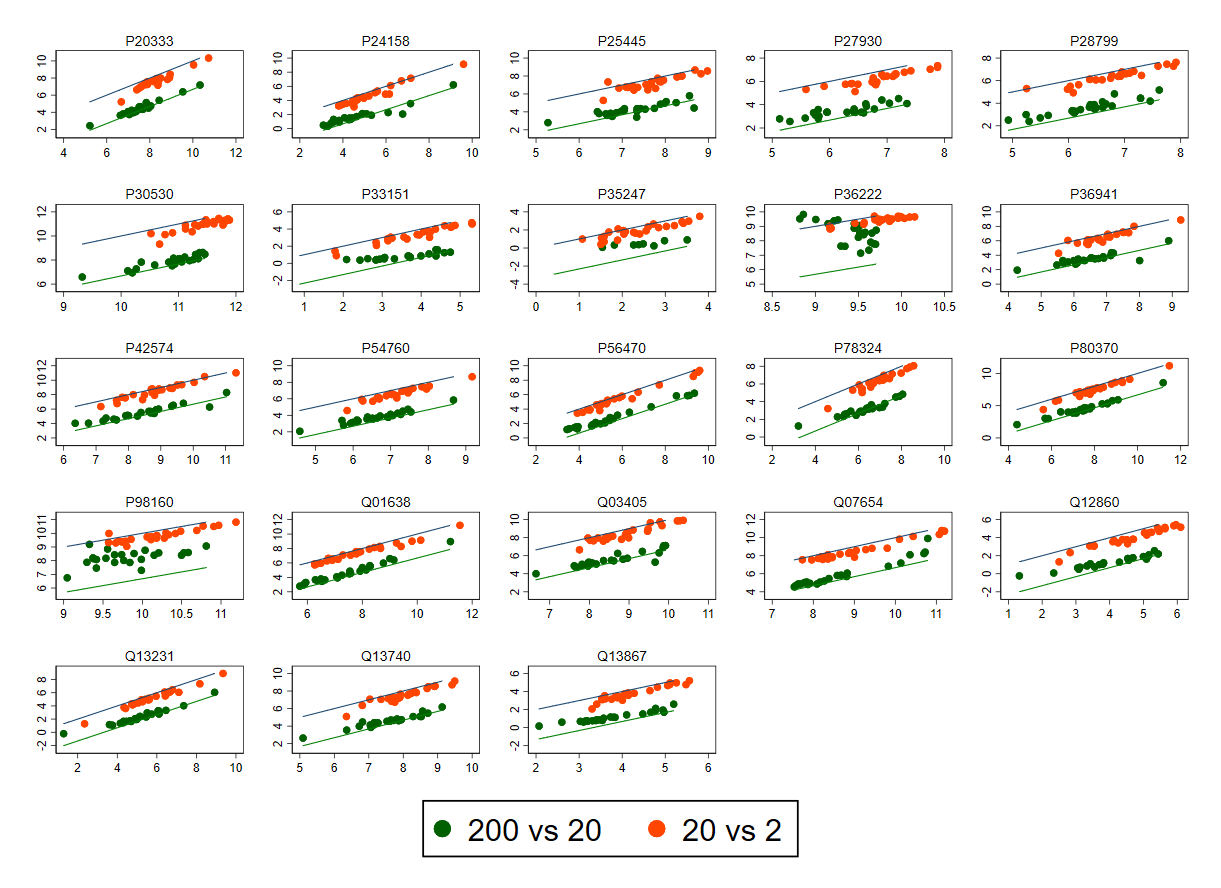

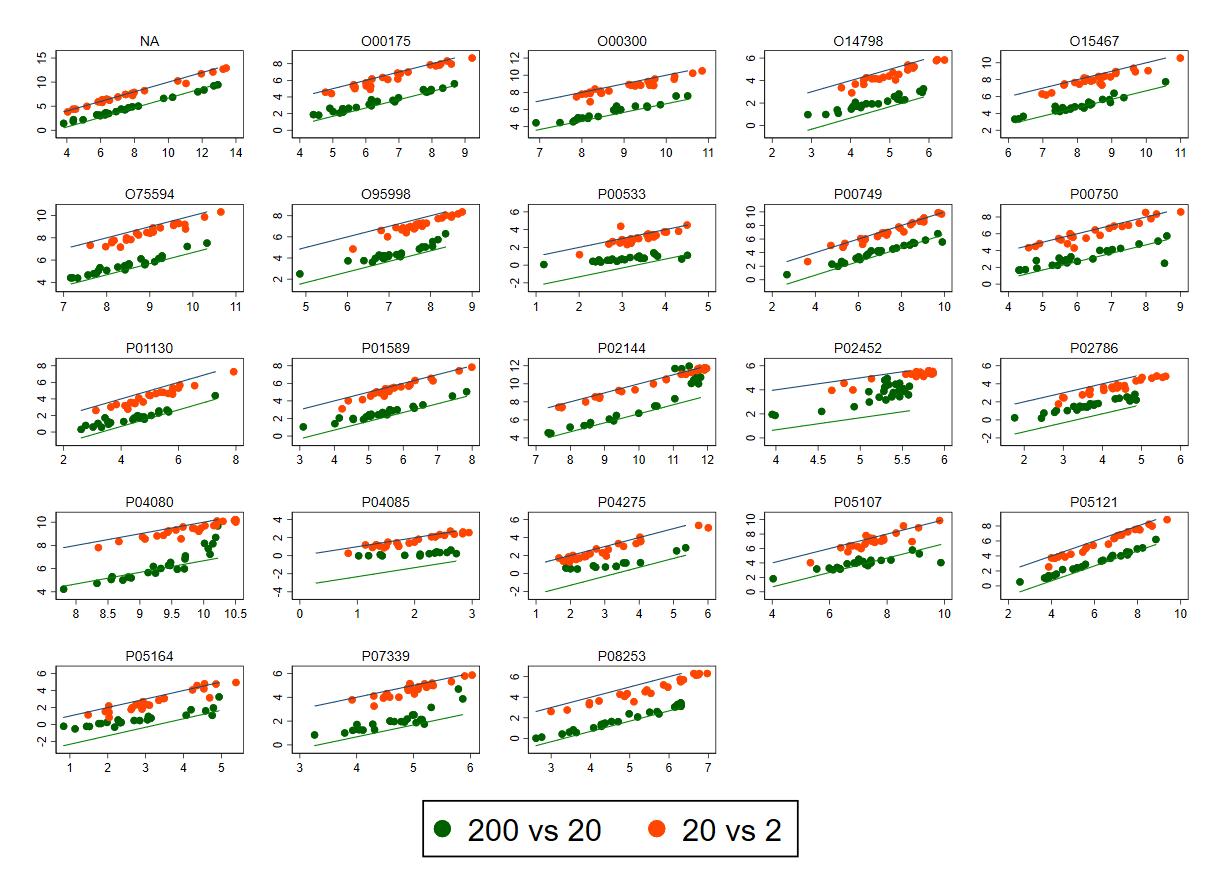

Supplementary figure 2. Dilution plots for all proteins in the CVDIII panel. The plots display the correlation between the different dilutions.

**Choice of dilutions**

The LOD is the lower limit and the ULOQ is the upper limit for the linear part of the standard curve. A standard curve for each protein is provided at the Olink webpage, but the equation is not stated, and hence the curve cannot be used. The provided curve has the concentration on the x-axis and NPX on the y-axis. The curve cannot be used to calculate the concentration, it can however be used to give an estimate of the linear range of the standard curve. Hence, it can be used to evaluate which dilution to use. The LOD (NPX) is measured each run, while the ULOQ (pg/ml) as well as the LOD (pg/ml) are provided at the Olink webpage for each protein. These values, as well as the relationship that a 1 NPX increase equals a doubling of the concentration, are used to calculate the equation for the linear part of the standard curve. The mean value of the LOD and the ULOQ provides the midpoint of this linear range. More than one dilution can provide a measured value within the LOD to ULOQ range, and to be able to choose one of the dilutions over the other, the dilution that gives a NPX that is closest to the midpoint was considered to be the best dilution. This was done for every sample for each protein, respectively. Lastly, the best dilution for a certain protein was determined by counting the number of samples with a certain dilution giving the NPX closest to the midpoint, and then choosing the most common dilution to use for all samples. For some proteins, the appearance of the dilution plot helped the determine which dilution was the best to use.

COMP was excluded from the differential expression analysis due to anomalies in its dilution plot (see Methods). However, it is one of the more established potential biomarkers for OA. Hence, the differential expression of COMP in the OA-group relative to the healthy group was calculated separately, see Supplementary Figure 3. A dilution of 1:4000 was used, since it is likely that the assay was saturated at lower dilutions. However, results were missing for four samples (two healthy and two OA) at this dilution. The mean difference in the NPX-values between the OA samples and the healthy samples, after being adjusted for BMI, sex and age, was -0.42, 95% CI [-1.05, 0.22]. Since the CI for COMP includes zero, we consider it less interesting to look at further in this study.

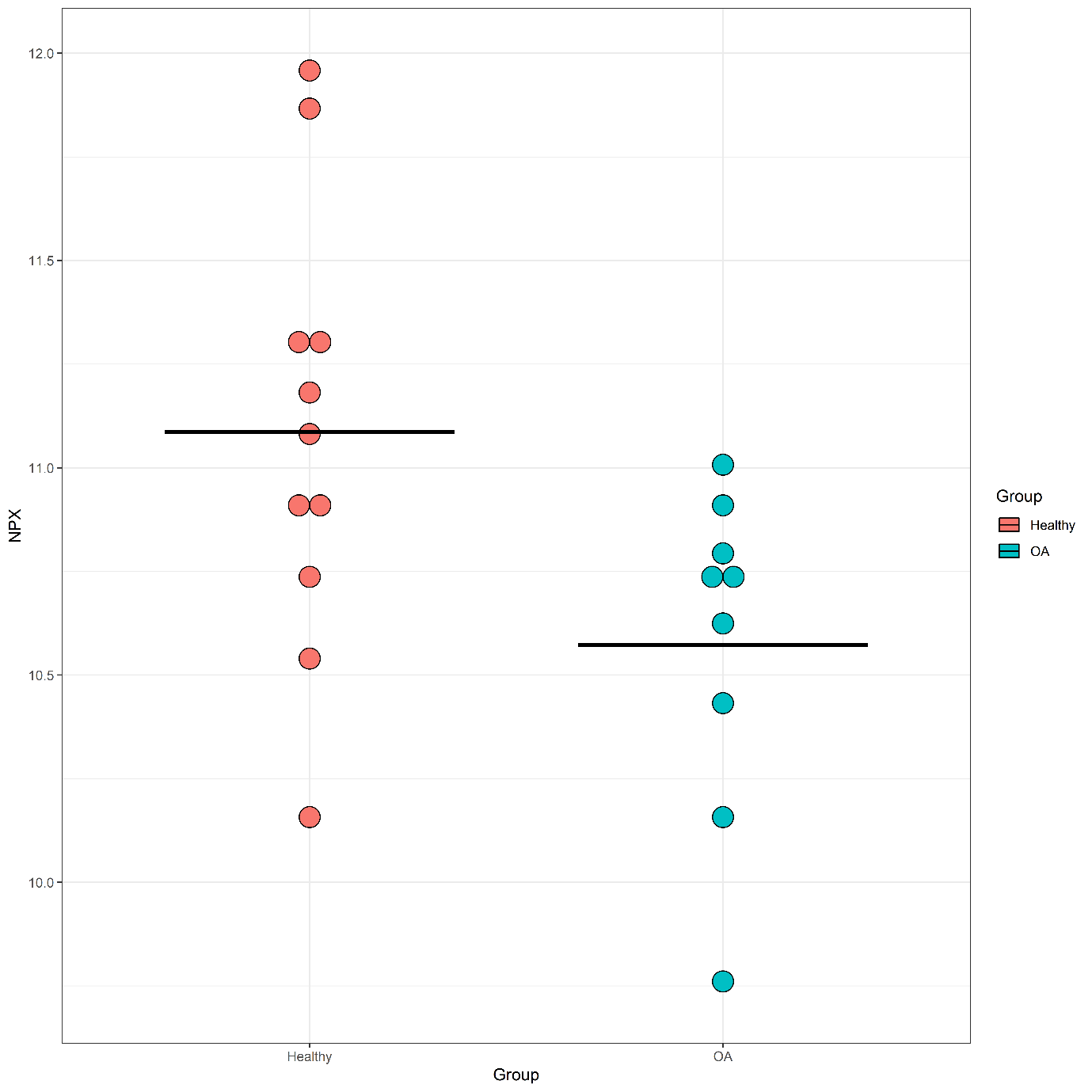

Supplementary figure 3. A groupwise comparison of the NPX-values for COMP in the individual samples. 11 healthy samples and 9 OA samples were run with x4000 dilution. The black bar represents the average value for each group.

**Differential network analysis**

We have data on 274 proteins from 3 Olink panels in synovial fluid of 11 TKR patients and 13 healthy donors. One protein has 2 missing values and was excluded. We attempt to create gaussian graphical model based on partial correlations and to derive potentially differential networks in the two groups. We used Joint Graphical Lasso (<https://www.ncbi.nlm.nih.gov/pmc/articles/PMC4012833/>) fused approach.

We selected the best models based on information criteria, where we consider AIC, BIC and EBIC. For the EBIC we used gamma value of 0.75. We aim for networks that are simple enough to be interpretable but complicated enough to be interesting.

We evaluated the criteria across a grid of lambda1 and lambda2 values, in a sequential order:

a) First, we set lambda2 to a low constant (0.01) and evaluate a set of lambda 1 from relatively low (i.e. retaining relatively many proteins) to relative high (i.e. sparse). Lambda1: from 0.13 to 0.89 by step of 0.04.

AIC suggested a large model with many proteins (low lambda1), while BIC suggest a sparse model (high lambda1). Given that it is known that AIC often suggests too complicated models, we will continue with high lambda1.

b) Second run: lambda 1 from 0.8 to 0.89 by 0.01, lambda 2=0.01, 0.005, 0.001.
c) Third search is then for lambda 1 from 0.86 to 0.89 by 0.01 and lambda2=(0.0005, 0.0001, 0.0075, 0.005, 0.0025, 0.001, 0.01, 0.02, 0.03, 0.04, 0.05) to finetune lambda2.

It is worth noticing that there are many models that have somewhat similar values, and thus selecting a model based only in information criteria would be challenging. Further, many models are very similar to each other (ex differ in only one or two edges) and thus it is not very meaningful to consider them as separate models.

Based on EBIC values, we selected 3 models that could be acceptable. A simple network (lamba1=0.86, lambda2=0.03):

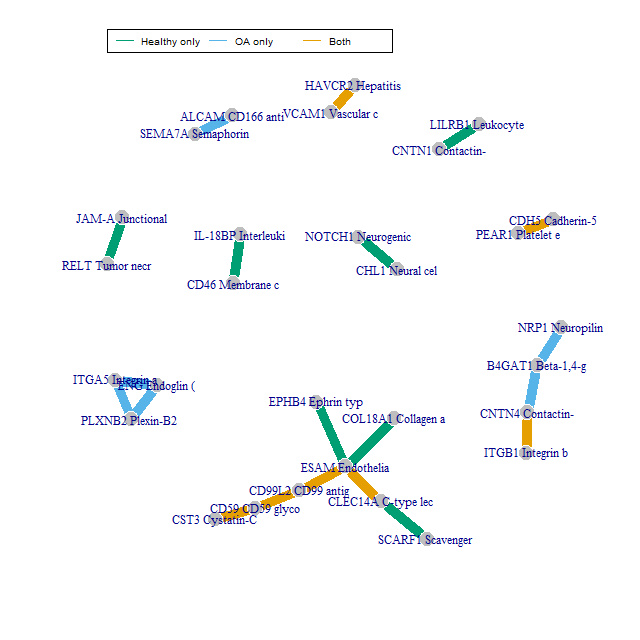

Moderate network (lambda1=0.87, lambda2=0.0075)

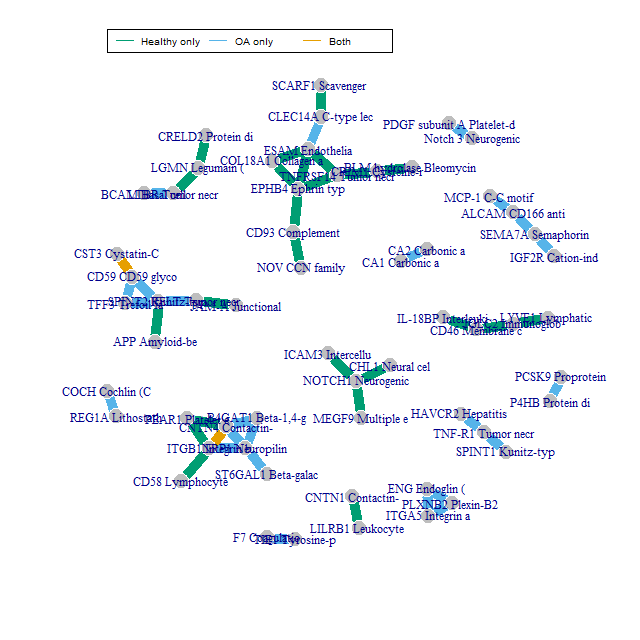

Large network (lambda1=0.86, lambda2=0.01)

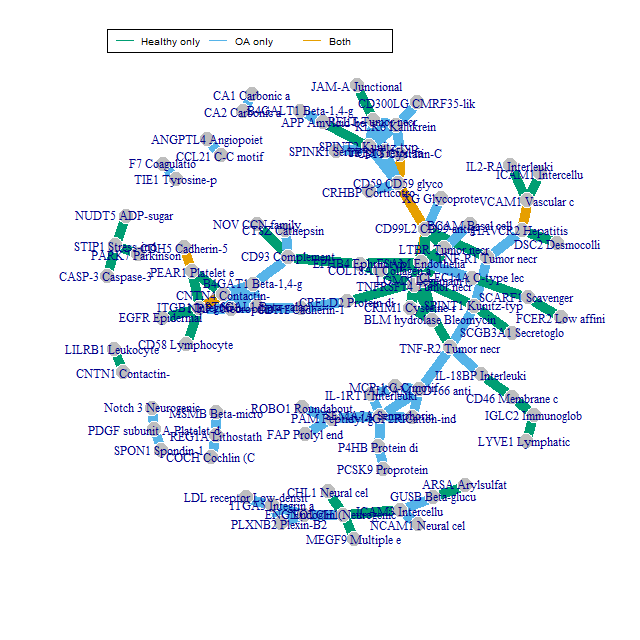

The models are almost nested within each other:

| First model | Small network | Moderate network |
| --- | --- | --- |
| Second model | Moderate network | Large network |
| Total number of proteins, first model | 29 | 58 |
| Total number of proteins, second model | 58 | 92 |
| Number of proteins only in first model | 3 | 0 |
| Number of proteins only in second model | 32 | 34 |
| Number of proteins only in both models | 26 | 58 |
| Number of edges only in first model | 4 | 0 |
| Number of edges only in second model | 34 | 59 |
| Number of edges only in both models | 16 | 50 |
| Number of proteins upregulated in TKR vs donors in differential expression analysis | 18 | 26 |
| Number of proteins downregulated in TKR vs donors in differential expression analysis | 6 | 11 |
